# Identification of two aptamers binding to *Legionella pneumophila* with high affinity and specificity

**DOI:** 10.1101/2019.12.13.875476

**Authors:** Mariam Saad, Deanna Chinerman, Maryam Tabrizian, Sebastien P. Faucher

## Abstract

*Legionella pneumophila* (*Lp*) is a water borne bacterium causing Legionnaires’ Disease (LD) in humans. Rapid detection of *Lp* in water system is essential to reduce the risk of LD outbreaks. The methods currently available require expert skills and are time intensive, thus delaying intervention. In situ detection of *Lp* by biosensor would allow rapid implementation of control strategies. To this end, a biorecognition element is required. Aptamers are considered promising biorecognition molecules for biosensing. Aptamers are short oligonucleotide sequence folding into a specific structure and are able to bind to specific molecules. Currently no aptamer and thus no aptamer-based technology exists for the detection of *Lp.* In this study, Systemic Evolution of Ligands through EXponential enrichment (SELEX) was used to identify aptamers binding specifically to *Lp*. Ten rounds of positive selection and two rounds of counter-selection against two *Pseudomonas* species were performed. Two aptamers binding strongly to *Lp* were identified with *K_D_* of 116 and 135 nM. Binding specificity of these two aptamers to *Lp* was confirmed by flow cytometry and fluorescence microscopy. Therefore, these two aptamers are promising biorecognition molecules for the detection of *Lp* in water systems.

## INTRODUCTION

*Legionella pneumophila (Lp)* is a pathogenic Gram-negative bacterium responsible for two types of respiratory diseases, namely the severe pneumonia Legionnaires’ Disease (LD) and the milder flu-like Pontiac fever ^1^. *Lp* occurs in both natural and engineered water systems and is one of the most prevalent pathogens in man-made, engineered water systems ^2^. Infections occur when the bacteria are aerosolized, and the contaminated aerosols are inhaled; *Lp* can then infect and replicate inside alveolar macrophages ^3^. Modern water systems provide optimal transmission conditions for *Lp* by generating aerosols ^4^. Leading sources of infection are cooling towers, hot water distribution systems, humidifiers, misters, showers, fountains, spa pools and evaporative condensers ^5^.

Outbreaks of LD consistently occur globally and have increased in recent years. The average incidence rate is about 10-15 cases per million people ^6^. According to the Centre for Disease Control, incidences of legionellosis have increased by four and a half times between 2000 and 2016 ^7^. The rise in LD outbreaks can be attributed to several factors such as aging infrastructures and an aging population who is more vulnerable to such infections, as well as increases in diagnosis and reporting ^8,9^. Nevertheless, most LD outbreaks are the consequences of management failure of man-made water systems ^10^. Examples of these failures for water distribution systems include keeping the temperature of hot water distribution system below 50°C and allowing water to stagnate ^10^. For cooling towers, lack of regular cleaning and disinfection is associated with increased risk ^10^. In both cases, routine monitoring of *Lp* is critical to evaluate risk, initiate treatment of water systems, and prevent outbreaks ^10^. The European Center of Disease Control (ECDC) specifies that immediate corrective measures should be taken when *Lp* levels reach a value of 10,000 CFU/L ^11^.

Currently, there are two ISO-certified strategies to detect *Lp* from water systems: the standard plate count method (AFNOR NF T90-431, ISO 11731) and qPCR (AFNOR NF T90-471, ISO/TS 12869). The plate count method is the gold standard for detecting *Lp* and involves its cultivation on selective media and the enumeration of bacterial colonies showing *Lp*-specific morphology ^12,13,14^. The whole procedure takes up to 14 days which delays the application of disinfectant and increase the chances of outbreak ^15^. A pilot study performed in 2011 evaluated the consistency of the results obtained by this method between several different laboratories. Qualitative results did not differ drastically between laboratories, but quantitative results showed large variation, within and between laboratories ^16^. Therefore, the culture method should be use with caution to precisely enumerate *Lp.* A second major limitation is the presence of viable but non-culturable (VBNC) *Lp* cells which leads to an underestimation of the true amount of infectious *Lp* in a system ^17,18^. The qPCR method relies on the quantification of *Legionella* DNA. Its major advantages in comparison with conventional culture method is the rapid turn-around time, high sensitivity and specificity, low limit of detection, as well as the ability to detect VBNC cells. When used in conjunction with the culture method, qPCR can serve as a powerful tool. There are, however, several drawbacks: qPCR typically overestimates *Lp* burden because it detects dead cells ^19,20^ and the presence of PCR inhibitors may prevent the use of this method. In addition, multiple processing steps are required which increases the overall cost of the qPCR method ^21^. Unfortunately, it is impossible to develop these two methods into rapid, cost-effective, sensitive tests that would detect *Lp* in real-time, on-site, without any additional processing steps ^22,23^.

Biosensors are attractive detection technology that could address the problems associated with culture-based bacterial detection methods. These analytical devices are commonly used to assess and quantify in real-time and with high sensitivity the presence of an analyte such as a protein, peptide or cell in a fluid ^24^. However, a biosensing approach to *Lp* detection would require a specific biorecognition element, which when coupled with a transducer, translates its interaction with *Lp* cells into a meaningful readout ^24^.

Various biorecognition elements, such as antibodies, lectins or aptamers, can be used. The latter are becoming the primary choice for biosensing strategies due to their easily modifiable nature and versatility ^25,26^. Aptamers are antibody analogues. They are short single stranded DNA or RNA oligonucleotides that can be cost effectively synthesized in a high throughput manner. The aptamer folds into a specific, stable structure and interacts with its targets via shape complementarity, hydrogen bonding, electrostatic interactions and stacking interactions ^27^. This allows its binding with high affinity and specificity to a wide variety of targets ranging from small molecules, peptides, proteins to whole cells ^27^. A key characteristic of aptamers is the possibility to generate them *in vitro* in the same condition as those used for detecting the analyte. This is a clear advantage over antibodies which are produced under strict physiological conditions ^28^. In addition, aptamers can be easily modified and therefore be optimized for various sensing platforms such as lateral flow assays, surface plasmon resonance sensors, flow cytometry or fluorescence microscopy ^29^.

The procedure by which an aptamer is created is known as Systemic Evolution of Ligands through EXponential enrichment (SELEX). Conceived in 1990 by the teams of Gold and Szostak ^30,31^, SELEX is an iterative process which involves incubating a target with a large library of oligonucleotides, separating the target bound and unbound oligonucleotides and then amplifying the target bound sequences via PCR for the next round of selection. The selection rounds are repeated until the oligonucleotide pool is enriched with sequences that bind specifically and with high affinity to the target ^31^. Over the years, many variations of the original SELEX methods were published. Among those, one is particularly useful for the present study. Cell-SELEX can be used to select aptamers binding to whole living cells and, thus, eliminates the need for prior knowledge of a target molecule ^32^. Rounds of counter-selection are typically used to reduce aptamer cross-reactivity across targets by eliminating non-specific aptamers ^29^. Cell-SELEX has been successfully employed to isolate aptamers against various bacterial species such as *E. coli, Salmonella typhimurium, Campylobacter jejuni, Listeria monocytogenes, Staphylococcus aureus* and *Vibrio parahaemolyticus* ^28, 33–37^. Several of these aptamers are used in conjunction with optical, mechanical or electrical/electrochemical biosensors to mitigate the problems associated with traditional bacterial detection methods. Although numerous works have been done to detect *Lp* with the use of biosensors ^38–40^ no study has used or even created aptamers binding to *Lp*. Consequently, no aptamer and thus no aptamer-based technology currently exists for the detection of *Lp.*

In this work, the cell-SELEX procedure was employed to generate aptamers against *Lp*. Two *Pseudomonas* species were used for counter-selection to improve the specificity of the aptamers. Two aptamers were identified and their binding affinity and specificity for *Lp* were evaluated by flow cytometry and fluorescence microscopy.

## MATERIALS AND METHODS

### Bacterial Strains and Culture Conditions

The environmental *Lp* strain *lp120292*, isolated from a cooling tower implicated in the 2012 outbreak in Quebec City, was used as the target strain for aptamer generation (Levesque 2016). The strain Lp*GFP is *lp120292* transformed with plasmid pXDC31 expressing the green fluorescent protein (GFP) under the *Ptac* promoter ^41^. The thymidine auxotroph *Lp* strain Lp02, derived from *Lp* Philadelphia-1 was used to confirm binding of the aptamers ^42^. *Lp* was cultured on CYE (ACES-buffered charcoal yeast extract) agar plates supplemented with 0.25 mg/ml L-cysteine and 0.4 mg/ml ferric pyrophosphate, at 37 °C for 3 days. Lp*GFP strain was grown on CYE media supplemented with 5 μg/ml chloramphenicol and 1 mM isopropyl β-D-1-thiogalactopyranoside. For liquid culture, *Lp* was suspended in AYE (ACES-buffered yeast extract) broth supplemented with 0.25 mg/ml L-cysteine and 0.4 mg/ml ferric pyrophosphate until post-exponential phase (OD_600_ of 2.5). *Pseudomonas putida* KT2440 and *Pseudomonas fluorescens* LMG1794, were first cultured on LB agar plates at 30 °C for 24 hours and then grown in LB medium (Difco) until the cultures reached post-exponential phase (OD_600_ of 2.0). *Pseudomonas sp., Brevundiomonas sp., Bacillus sp., Staphlyococcus sp.* and *Sphingomonas sp.* were isolated from cooling towers as part of a different study (Paranjape, in preparation). Briefly, the bacteria were isolated on nutrient agar incubate at 30 °C. Isolates were the classified using 16S rDNA sequencing. These strains were first cultured on nutrient agar plates (Difco) at 30 °C for 24 hours and then grown in nutrient broth medium (Difco) overnight until the cultures reached post-exponential phase (OD_600_ of 2.0-2.5).

### Oligonucleotides Library and Primers

A random ssDNA library of 10^15^ sequences was chemically synthesized and purified by HPLC (Integrated DNA Technologies). The library consisted of a central random region of 45 nucleotides flanked by two different primer binding regions at the 5’ and 3’ ends: 5’-GCAATGGTACGGTACTTCC-45N-CAAAAGTGCACGCTACTTTGCTAA-3’. The forward and reverse primers were conjugated with fluorescein (FITC) and biotin, respectively. The forward primer (FP) sequence is 5’-fluorescein-GCAATGGTACGGTACTTCC-3’. The reverse primer (RP) sequence is 5’-biotin-TTAGCAAAGTAGCGTGCACTTTTG-3’ ^28^. FITC was used to quantify ssDNA and monitor the SELEX procedure via flow cytometry. The biotin was used in conjunction with streptavidin coated magnetic beads (Promega) to generate ssDNA from the amplified double-stranded aptamer pool following PCR. For cloning and sequencing, PCR was performed with unmodified versions of the primers.

### Bacterial Cell-SELEX procedure

Cell-SELEX was performed as previously described ^28^. Each round of SELEX consisted of three steps: Binding and elution, amplification, and recovery of ssDNA. Ten rounds of positive selection and two rounds of counter selection were performed (Figure 1). The first round of counter selection was performed with *P. putida* KT2440; the second one with *P. fluorescens* LMG1794.

**Figure 1:**
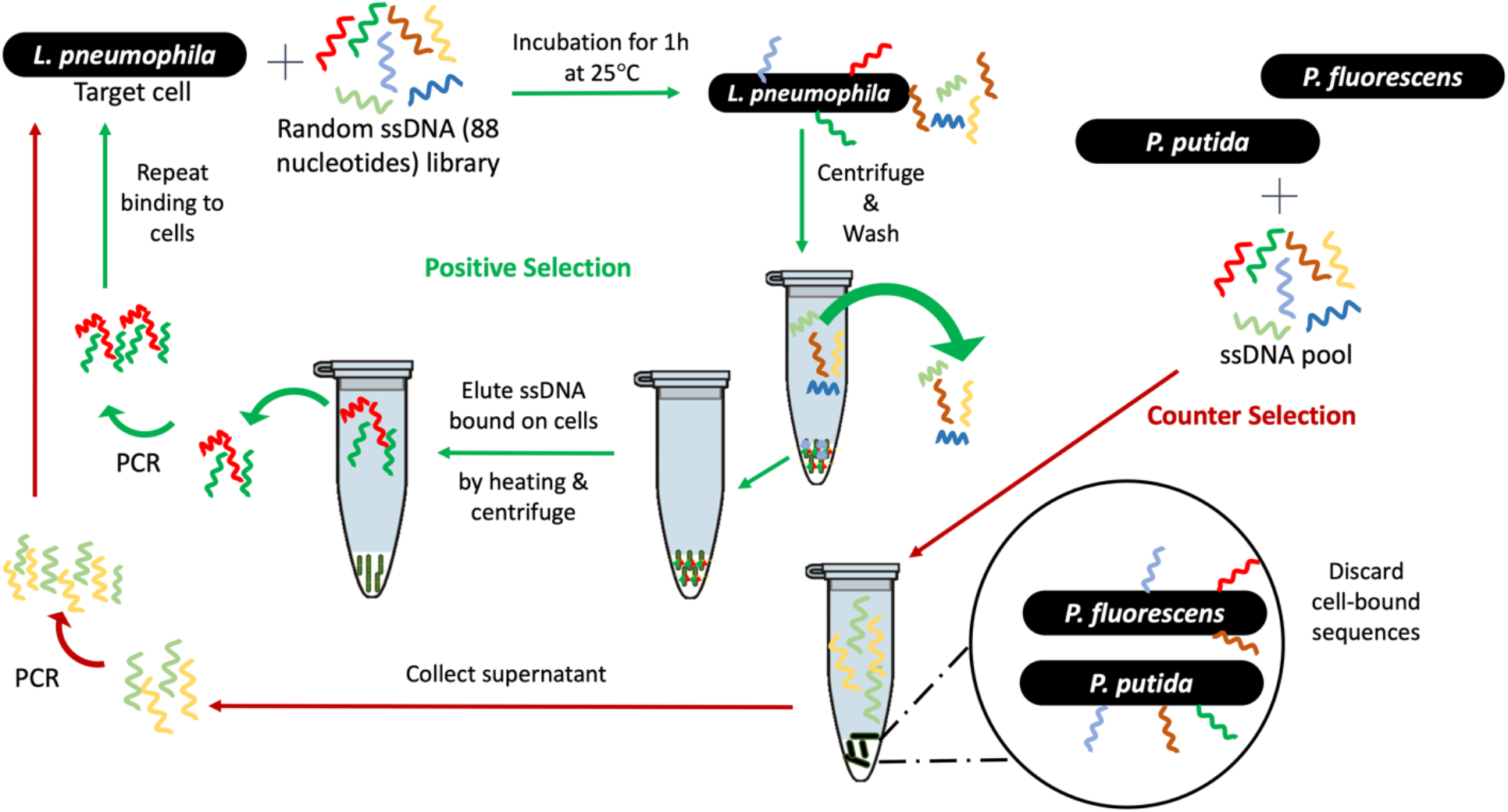
Schematic illustration of bacterial cell-SELEX procedure used in this study. A random library of oligonucleotides is incubated with *Lp lp120292* at room temperature for 1h. Sequences that do not bind are washed off and the cell-bound sequences are then released and amplified via PCR. The resulting sequences are then submitted to another round of positive selection. *P. fluorescens* and *P. putida* were used to perform counter-selection rounds to eliminate non *Lp*-specific sequences.

#### Binding and Elution

Cell-SELEX was performed with cells suspended in an artificial freshwater medium (Fraquil) to replicate the physiological state of nutrient-limited environmental conditions ^43,44^. Fraquil was prepared as described previously ^45^ with a final iron concentration of 10 nM and filter-sterilized using a 0.2 um filter (Sarstedt). Post-exponential phase cultures were rinsed twice with Fraquil (6000 *g*, 15 minutes) and suspended in Fraquil at an OD_600_ of 1 corresponding to a concentration of 10^9^ CFU/ml. The concentration of cells was confirmed by CFU counts for each round. The suspension was incubated at room temperature for 24 h. Fraquil exposed cells were washed three times in 1X binding buffer (phosphate buffered saline with 0.1 mg/ml salmon sperm DNA, 1% bovine serum albumin, and 0.05% Tween 20) at room temperature (25°C) using 6,000 *g* for ten minutes. The cell pellets were then suspended in 330 μl of 1X binding buffer. The aptamer pool was denatured by heating at 95 °C for 10 minutes, cooled immediately on ice for 10 minutes, and added to the cell suspension. Finally, 1X binding buffer was added to a total volume of 1 ml. For the first round, 32 μg of the initial library was used. For the subsequent rounds, approximately 400 ng of aptamer pool was used. The final mixture was incubated at 25 °C for 1 hour with mild shaking using a tube rotator at 150 rpm. Following incubation, the mixture was centrifuged at 6000 *g* for 10 minutes and washed twice with wash buffer (phosphate buffered saline containing 0.05% Tween 20) to remove unbound sequences. To elute the bound sequences from the cells, the final cell pellet was resuspended in 100 μl nuclease free water (Ambion) and heated at 95 °C for 10 minutes and immediately placed on ice for 10 minutes. After centrifuging at 6,000 *g* for 10 minutes at 25°C, the supernatant was collected and purified using overnight ethanol precipitation at −20 °C with 5 μg of glycogen as a carrier to recover the eluted ssDNA. The pellet was recovered, dried and suspended in nuclease free water (Ambion). The concentration and quality of the ssDNA was determined using a Nanodrop spectrophotometer (Thermofisher). For counter-selection the supernatant containing the unbound sequences was collected and purified via ethanol precipitation, as described above. To ensure there was no amplification or collection of unwanted bacterial DNA (instead of the desired amplification and collection of ssDNA oligonucleotides), a control sample consisting of bacterial cells without aptamer was included in each round.

#### PCR amplification

The purified aptamer pool was then amplified by PCR with One Taq DNA polymerase (NEB), according to the manufacturer’s protocol. All primers were used at a final concentration of 0.5 μM. PCR conditions were as follows: initial heat activation at 95 °C for 5 min and 25 cycles of 95 °C for 30 s, 56.3 °C for 30 s, 72 °C for 10 s, and a final extension step of 10 min at 72 °C. After amplification, the concentration and size of the PCR product were confirmed by gel electrophoresis using a 2.0% agarose gel. PCR products were then purified using a MinElute PCR Purification Kit (Qiagen). As expected, no amplification was observed for the control samples, lacking aptamer template.

#### Recovery of ssDNA

Streptavidin coated magnetic beads (Promega Technology) were used, according to the manufacturer’s recommendation. Briefly, 600 μg of magnetic beads were washed twice and then resuspended in 900 μl of washing buffer (phosphate buffered saline with 0.05% Tween 20). Next, approximately1 μg of PCR product was incubated with the magnetic beads for 10 min, mixing gently by inversion after every few minutes. The mixture was then washed in 1 ml of washing buffer. Finally, the beads were incubated with 500 μl of 200 mM NaOH for 5 minutes. The supernatant was then collected, and the FITC-labelled ssDNA was purified using ethanol precipitation as mentioned previously and quantified with a Nanodrop spectrophotometer (Thermofisher).

### Monitoring of SELEX by flow cytometry

The binding of the FITC-labelled aptamer pools from rounds 1 (R1), 6 (R6), 7 (R7), 8 (R8) and 10 (R10) to *Lp* was assessed using flow cytometry. Briefly, 35 nM of aptamer pools from each of these rounds was incubated with 10^6^ CFU/ml of *Lp* cells at 25 °C for 1 hour. Analysis was performed on a Guava easyCyte (Millipore) using the green fluorescence channel. A total of 5,000 events were recorded. Unlabeled cells were used as a control to measure autofluorescence. The Lp*GFP strain, producing strong green fluorescence from GFP, was used to adjust the gain of the green fluorescence channel. For analysis, a gate was first defined based on the forward and side scatters that included most of the cells. Then, a histogram of the number of cells vs the fluorescence intensity was used to define a region named Green_Lp where cells were considered positive for green fluorescence and therefore stained with aptamers. This region was setup to include very few cells of the unstained control and therefore represent fluorescence above the autofluorescence. Aptamer pools from R10 alone, without cells, was also analyzed to ensure that the aptamer alone was not forming aggregates that would be confused with cells.

### Cloning and Sequencing

To identify sequences binding to *Lp*, the aptamer pool from the 10^th^ round of SELEX was cloned with the pGEMT-easy Cloning and Ligation Kit (Promega). To investigate the effect of counter-selection on the aptamer pools, we also cloned and sequenced aptamers from the 6^th^ round of SELEX. Positive colonies, containing aptamer inserts, were determined via blue-white screening and confirmed by PCR. Plasmids were extracted and purified using a Miniprep Kit (Qiagen) and sequenced by Sanger Sequencing at the Plate-forme d’Analyse Génomique of Laval University. Secondary structures of the aptamer sequences were determined using the Mfold web server using default parameters ^46^.

### Characterization of aptamers R10C5 and R10C1

The binding of the aptamers R10C5 and R10C1 to *Lp* and to the species used for counter-selection was characterize further. R10C5 and R10C1 were individually synthesized with FITC at the 5’ end (Integrated DNA Techology).

#### Determination of the Disassociation constant (*K_D_*)

To determine the *K_D_* of R10C5 and R10C1, varying concentrations of FITC-tagged aptamers (1000 nM, 100 nM, 10 nM, and 1 nM) were incubated with 10^6^ CFU/ml of *Lp* cells suspended in Fraquil and the fluorescence obtained at each concentration was measured using flow cytometry, as described above, in triplicate. The number of bound cells (FITC-positive) were recorded and used to determine the *K_D_* by interpolating the logarithmic curve using GraphPad Prism 7.03.

#### Specificity assay

To determine the specificity of R10C5 and R10C1 for *Lp* cells, the binding to counter-SELEX *Pseudomonas* strains as well as cooling tower isolates was tested using flow cytometry. All cells were suspended in Fraquil and prepared as described above for cell-SELEX. Briefly, 100 nM of R10C5 and R10C1 was incubated with 10^7^ CFU/ml of the strain used for SELEX (*lp120292*), another *Lp* strain (Lp02), the strains used for counter-selection (*P. putida KT2440* and *P. fluorescens LMG1794*), and the isolates from cooling towers (*Pseudomonas sp., Brevundiomonas sp., Bacillus sp., Staphlyococcus sp.* and *Sphingomonas sp.*) for 1 hour at 25 °C with mild shaking. The suspension was centrifuged for 10 minutes at 6000g to eliminate excess aptamer and resuspended in Fraquil. These suspensions were then analyzed using flow cytometry as described above. This experiment was done in triplicate. Cells suspended in Fraquil without any aptamer added were used as controls. The percentage of bound cells was determined as described above. Statistical differences were assessed using a one-way ANOVA and Dunnett correction for multiple comparison using Graphpad prism 7.03.

#### Fluorescence microscopy assay

FITC-labelled R10C5 and R10C1 aptamer (500 nM) were incubated with 10^8^ CFU/ml of target cell *lp120292* or counter-selection strain *P. fluorescens LMG1794* for 1 hour at 25 °C on a tube rotator at 150 rpm. Cells were suspended in Fraquil as mentioned previously. Negative controls included cells suspended in Fraquil without any aptamer and aptamer suspended in Fraquil, without cells added. The suspensions (10 μl) were dropped on a glass slide (Fisherbrand), and a #1.5 cover slip (Fisherbrand) was used to make a thin layer. Using a 63X oil immersion objective, brightfield and fluorescent images of bacteria were observed using an Apotome epifluorescence microscope (Zeiss) using pre-set excitation and emission filters for FITC, 495 nm and 519 nm respectively.

## RESULTS AND DISCUSSION

### Selection of Aptamers binding to *Lp*

Cell-SELEX was used to select aptamers binding specifically to *lp120292.* This strain was selected because it was involved in the Quebec City Outbreak in 2012 ^47^. To mimic the physiological state of *Lp* in water system, *Lp* cells were grown to post-exponential phase and suspended in Fraquil for 24 h at 25 °C to induce starvation and the associated morphological and physiological changes ^43,44^. Seven rounds of positive selection were performed, followed by one round of counter-selection, two rounds of positive selection, an additional round of counter-selection and a final round of positive selection (Figure 1). Two *Pseudomonas* strains were used for counter-selection because they are also Gram-negative proteobacteria frequently isolated from water systems where *Lp* is found.

To monitor the progress of the SELEX procedure and ensure that the proportion of sequences binding to *Lp* was increasing, the binding of the FITC-labelled aptamer pools from rounds 1 (R1), 6 (R6), 7 (R7), 8 (R8) and 10 (R10) to *Lp* was examined using flow cytometry. Cells incubated with the initial aptamer library showed minimal fluorescence compared to the negative controls (Figure 2, Lp Lib). Cells incubated with aptamer from the first positive selection round showed a drastic increase in fluorescence (Figure 2, Lp R1). The saturation in the fluorescence intensity and the percentage of bound cells starting at R6 suggests that the pool is dominated by sequences binding to *Lp* (Figure 2, R6, R7, R8, R10). A small decrease in fluorescence and percentage of bound cells at R8 suggests that the first counter-selection step removed a few sequences. The fluorescence intensity remains similar between round 8 and 10 indicating that the second round of counter-selection did not remove *Lp*-specific sequences and that our strategy was successful in retrieving aptamers binding to *Lp.*

**Figure 2:**
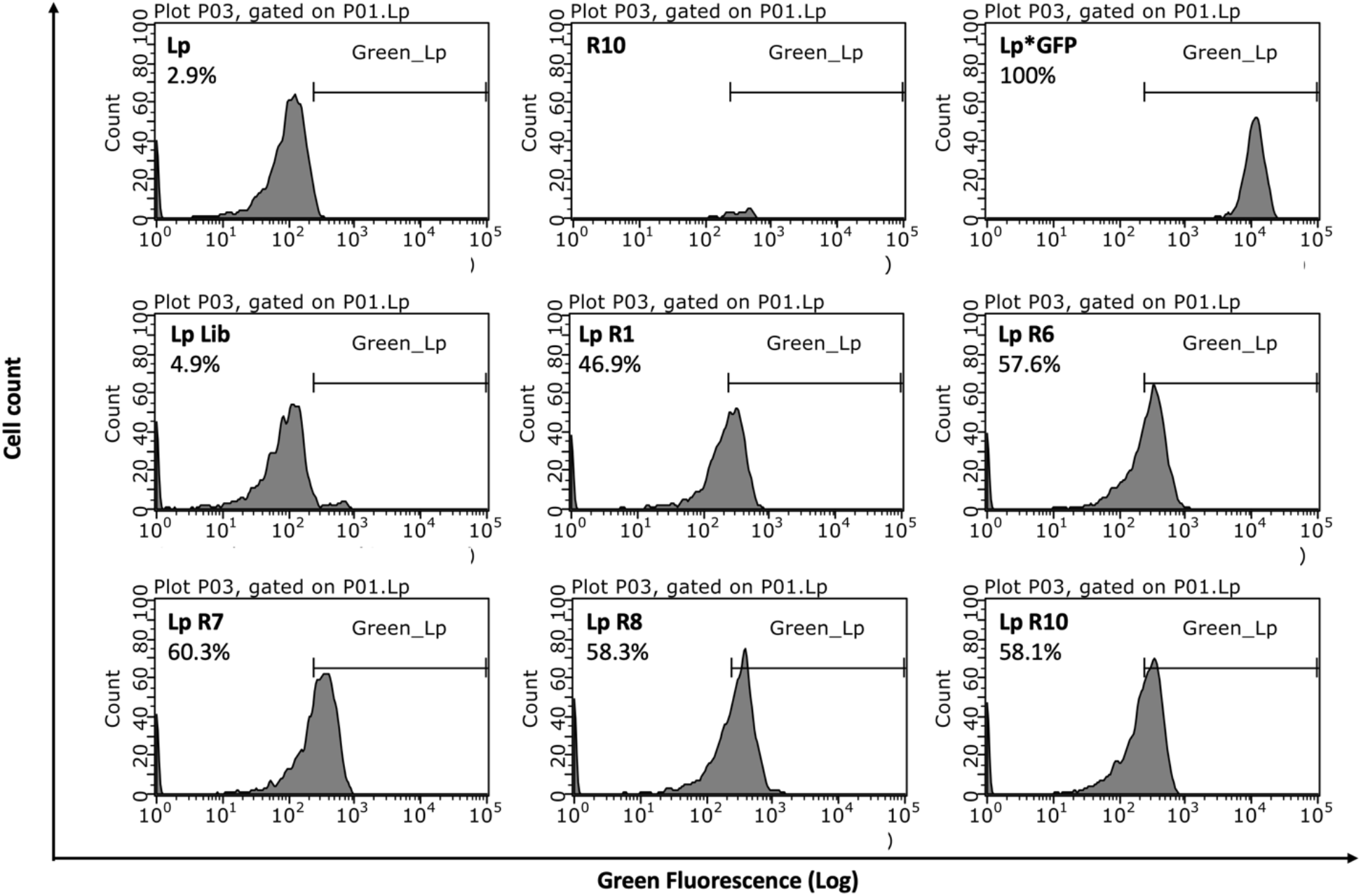
Fluorescent labeling of *Lp* with FITC-labeleld aptamers pools obtained after selected round of SELEX. *Lp* strain *1p120292* was incubate without aptamers (Lp), with 35 nM of the aptamer library (Lp-lib) and of the aptamer pools obtained after round 1 (Lp-R1), 6 (Lp-R6), 7 (Lp-R7), 8 (Lp-R8) and round 10 (Lp-R10). The fluorescence obtained with aptamer from round 10 alone (R10), without cells, was also evaluated. Lp*GFP is a GFP producing version of *lp120292* and is used as a positive control. The percentages refer to the proportion of cells with fluorescence above the autofluorescence, falling in the Green_Lp region.

### Cloning and Sequencing

Analyzing the sequences obtained from the 10^th^ round of positive selection allowed for identifying two different ssDNA aptamers, named R10C5 and R10C1 (Table 1 and Figure 3). Of the 13 sequences that were retrieved, 12 of them were R10C5 whereas 1 was R10C1. In contrast, the survey of a non-exhaustive list of the sequences present in the R6 aptamer pool revealed eight different sequences out of 9 clones, but none similar to R10C1 and R10C5. This illustrates the directional evolution of the pool as a result of the additional positive selection rounds and counter-selection steps ^48,49^. A strong bottleneck effect was likely caused by the last four positive selection rounds and the apparently stringent counter-selection rounds, which most likely led to the removal of several aptamers.

**Figure 3:**
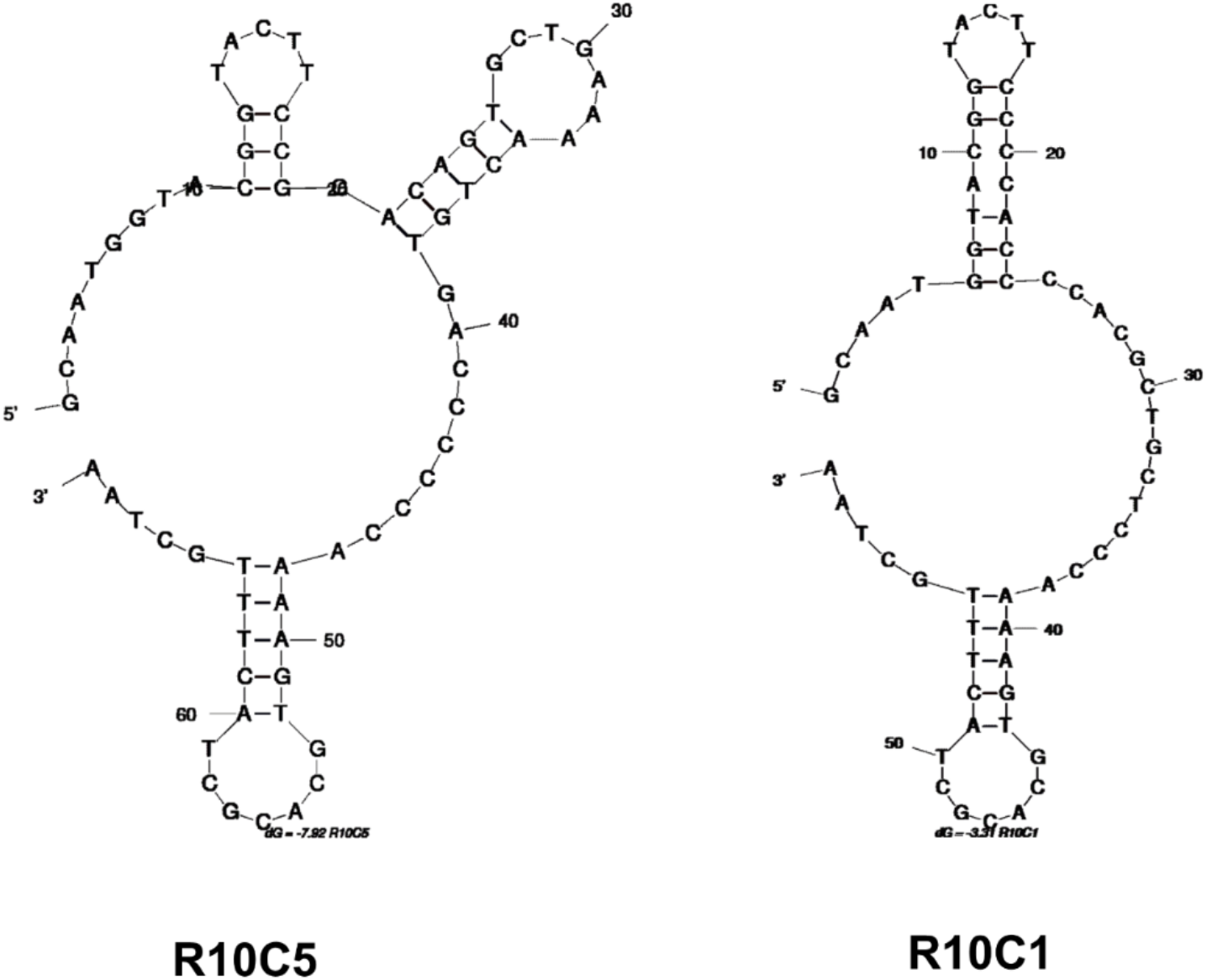
The structure of the aptamers R10C5 and R10C1 were determined using Mfold.

**Table 1:**
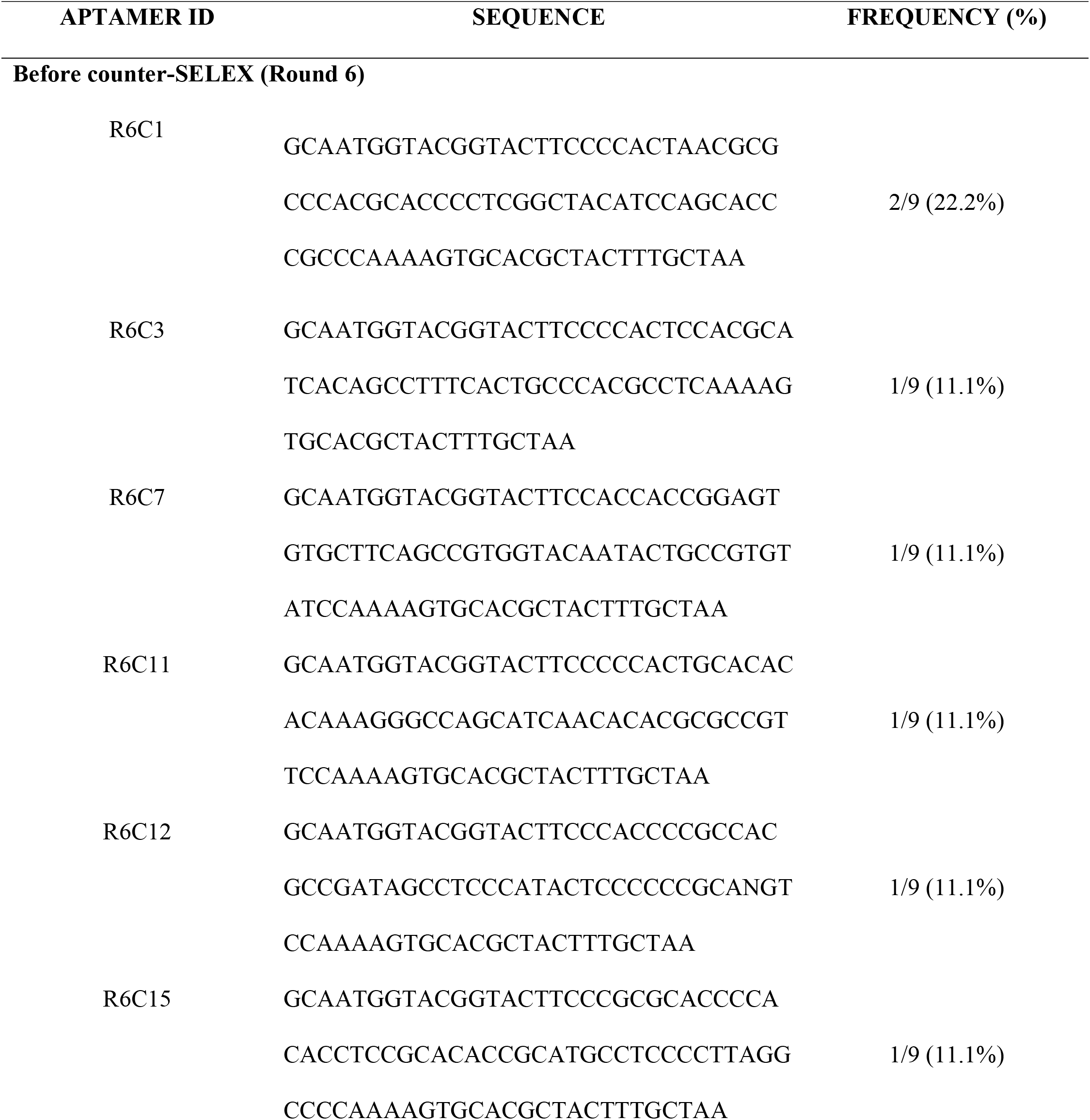
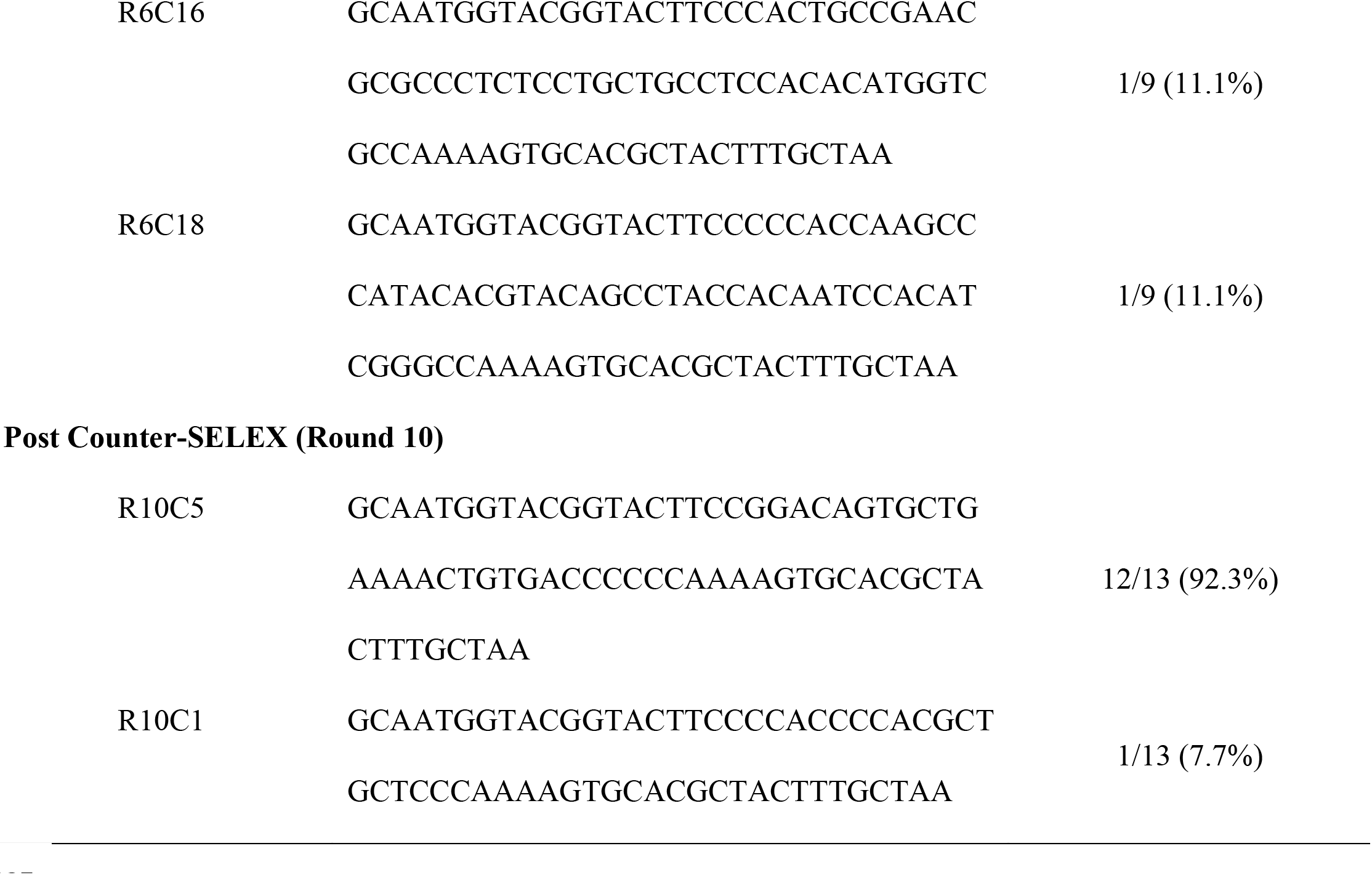
Aptamer’s sequences from round 6 and round 10.

### Determination of *K_D_*

The calculated *K_D_* is 116 nM for R10C5 and 135 nM for R10C1 (Figure 4). These values are comparable to high affinity antibodies that typically show nanomolar ranges of *K_D_* for small protein targets ^50^. These values are also consistent with published aptamers created against whole bacterial pathogens. For example, aptamers isolated against *Escherichia coli, Enterobacter aerogenes, Klebsiella pneumoniae, Citrobacter freundii, Bacillus subtilis*, and *Staphylococcus epidermidis* showed *K_D_* ranging from 9.22–38.5 nM ^51^. Two 62 nt aptamers binding to *Staphylococcus aureus* have *K_D_* of 35 nM and 129 nM ^52^.

**Figure 4:**
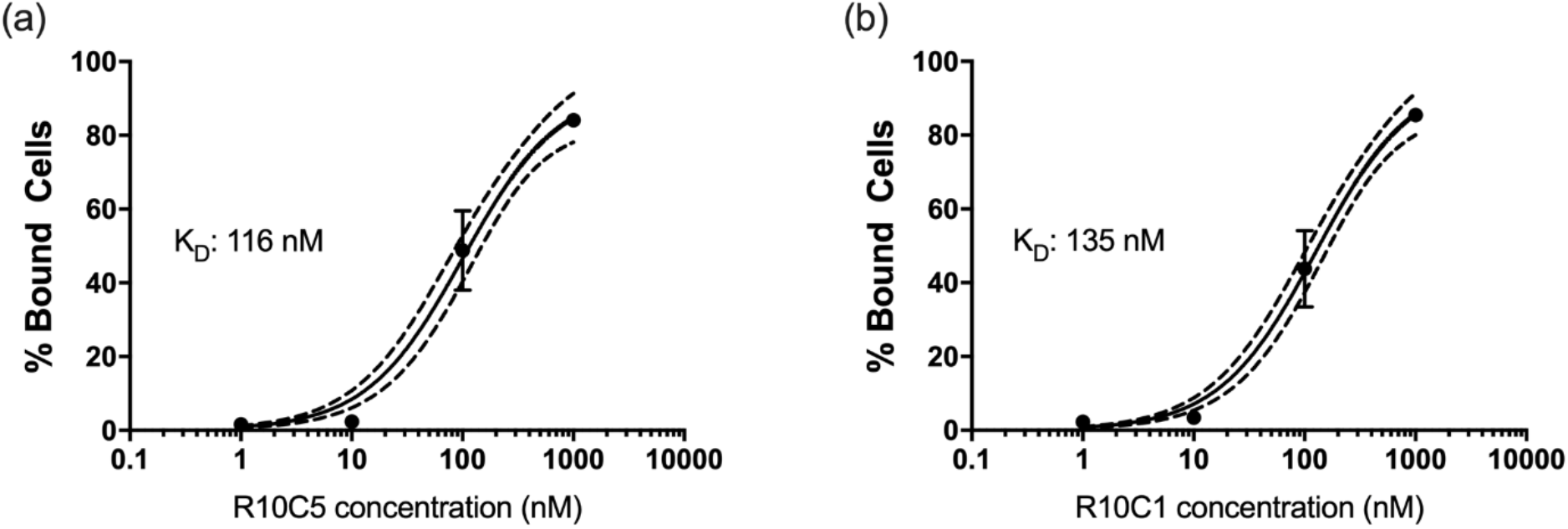
Determination of the *K_D_* of the aptamers R10C5 (a) and R10C1 (b). *Lp (lp120292)* was incubated with 10-fold dilutions of the FITC-tagged aptamers and the fluorescence was measured by flow cytometry. The number of cells displaying fluorescence above the autofluorescence were counted as bound cells, as described in Figure 2. The graphs show the average and standard deviation of three experiments. The equilibrium dissociation constant, *K_D_* was calculated using GraphPad Prism 7.03.

### Specificity of R10C5 and R10C1

Figure 5A shows the binding of R10C5 and R10C1 to the strains used for counter-selection. Around 60% of *lp120292* cells are stained by R10C5, consistent with previous results shown in Figure 2, but only 20% of *Pseudomonas* strains are labelled. Similarly, R10C1 shows significantly more binding to *Lp* than to *Pseudomonas* (Figure 5B). The aptamer R10C5 stained Lp02 similarly to *lp120292*, suggesting that the binding of this aptamer is not restricted to a particular strain of *Lp* (Figure 5A). Moreover, both aptamers show very low binding to environmental isolates from cooling tower water. In these cases, less then 10% of cells were labelled by the aptamers (Figure 5C and 5D). The specificity of these aptamers for *Lp* was further analyzed by fluorescence microscopy. Both aptamers strongly stained *Lp* (Figure 6) but not *P. fluorescens LMG1794*, one of the strains used for counter-selection. These results suggest that the aptamers R10C5 and R10C1 are highly specific to *Lp*.

**Figure 5:**
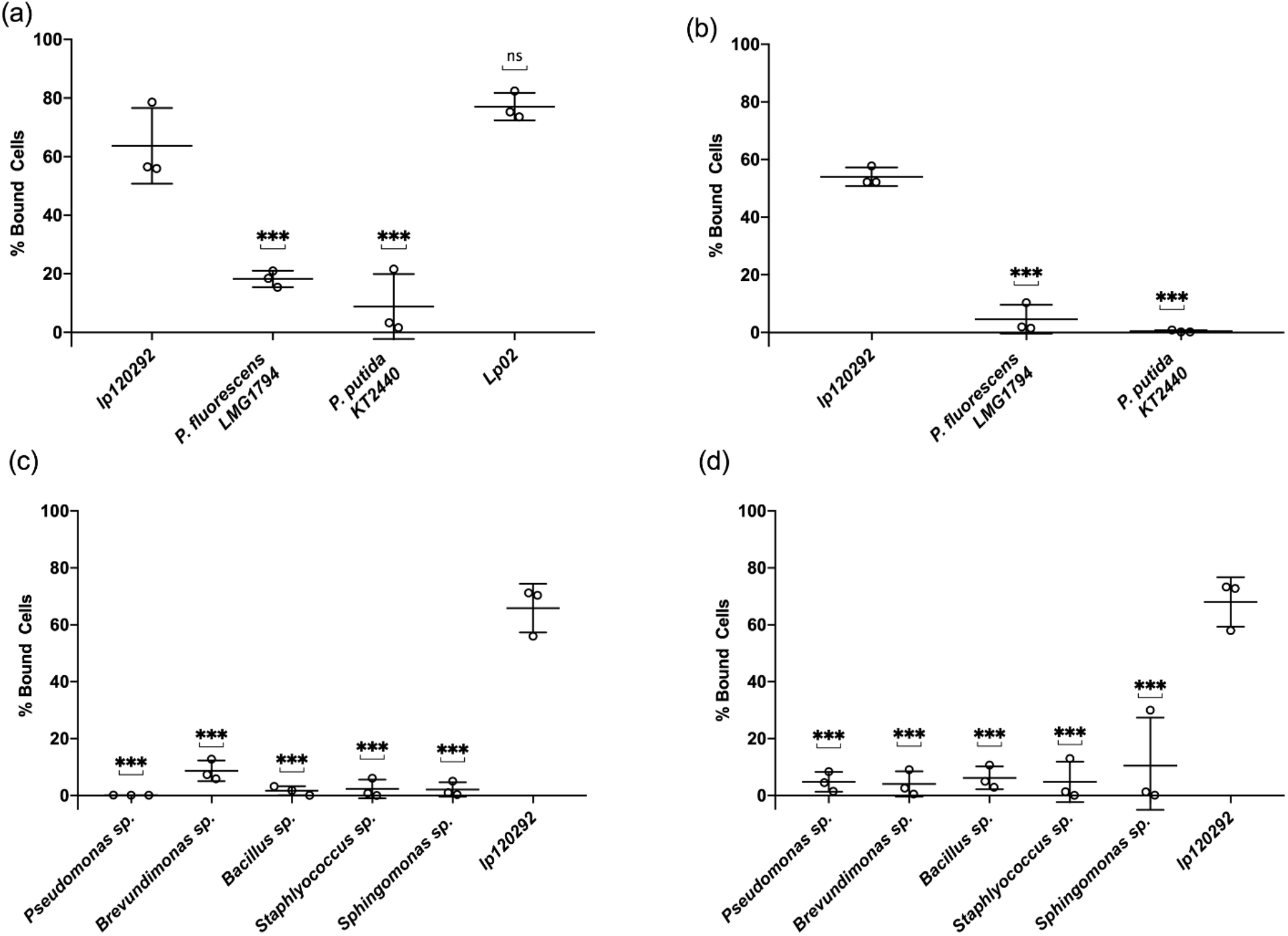
The specificity of FITC-labelled R10C5 and R10C1 aptamers binding to *Lp* strain *lp120292* (positive control), to counter SELEX *Pseudomonas* strains (a and b) as well as to environmental isolates (c and d) was analyzed by flow cytometry. The binding of R10C5 to *Lp* strain Lp02 was also analyzed. The percentage of cells bound by R10C5 (a and c) and R10C1 (b and d) to counter-SELEX strains (a and b) and environmental isolates (c and d) are presented. The values of three experiments are shown with the mean and standard deviation. A one-way ANOVA with a Dunnett correction for multiple comparisons was used to infer statistical significance compared to *Lp* strain *lp120292*: *** *P* < 0.001; ns, not significant.

**Figure 6:**
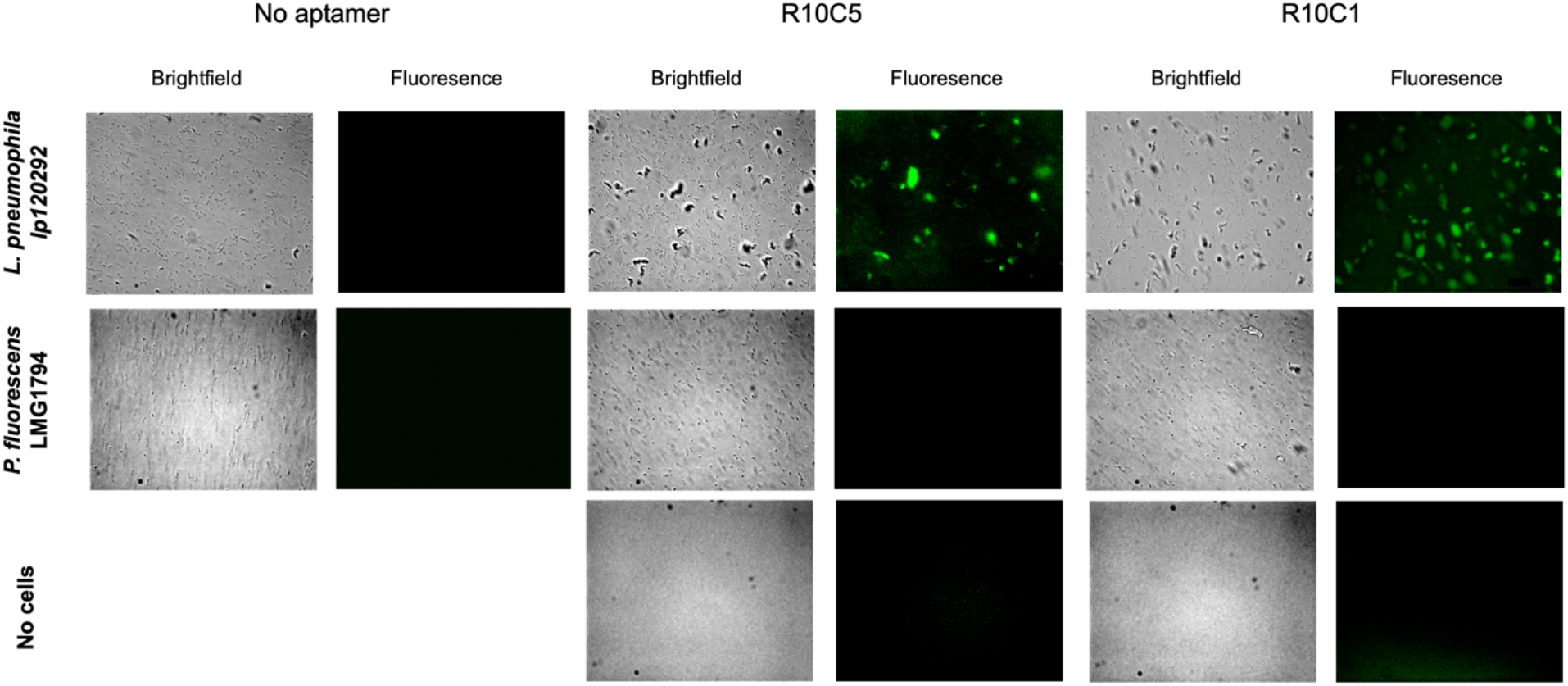
The specificity of FITC-labelled R10C5 and R10C1 aptamer was tested by measuring their ability to bind to *P. fluorescens* LMG1794 by fluorescence microscopy. Binding to *Lp* strains *lp120292* serves as a positive control. The no cells control consists of aptamer alone.

In conclusion, our cell-SELEX strategy was successful in identifying two aptamers binding to *Lp* with high affinity (*K*_D_ = 116 nM for R10C5 and 135 nM for R10C1). Whereas R10C5 seems to stain *Lp* more strongly then R10C1, the latter seems more specific to *Lp*, showing minimal binding to the counter SELEX *Pseudomonas* strain. Both aptamers showed minimal binding to cooling towers isolates, indicating that the aptamers are suitable to detect *Lp* in complex water samples. Modification of these aptamers could be attempted to further increase their affinity and specificity to *Lp*. Nonetheless, these aptamers are promising candidates as biorecognition elements to develop a biosensor to detect *Lp* in real time and *in situ.*

## ACKNOWLEDGEMENTS

We are grateful to Jonathan Perreault (INRS-IAF) for useful discussion and to Youssef Chebli (McGill University) for assistance with the fluorescence microscopes. The *Pseudomonas* strains used for counter selection are a kind gift from Eric Déziel (INRS-IAF). This study was supported by an NSERC Strategic Partnership Grant to MT and SPF. MS was supported by a CRIPA scholarship supported by the Fonds de recherche du Québec - Nature et technologies n°RS-170946.

## AUTHOR CONTRIBUTIONS STATEMENT

MS, MT and SPF designed the study. MS and SPF planned the experiments. MS and DC performed the experiments. MS wrote the first draft of the manuscript. MS, MT and SPF edited the manuscript. All authors approved the submitted version of the manuscript.

